# Hsa-MiR-483 -3p Regulates the Extracellular Matrix Proteins via TGFβ2/SMAD4 Signaling in the Glucocorticoid-responsive Human Trabecular Meshwork Cells

**DOI:** 10.1101/2025.01.14.633091

**Authors:** Ravinarayanan Haribalaganesh, Rajendrababu Sharmila, Ramasamy Krishnadas, Colin E. Willoughby, Srinivasan Senthilkumari

**Affiliations:** Department of Ocular Pharmacology, Aravind Medical Research Foundation, Madurai, Tamilnadu, India; Glaucoma Clinic, Aravind Eye Hospital, Madurai, Tamilnadu, India; Genomic Medicine, Biomedical Sciences Research Institute, Ulster University, Northern Ireland, United Kingdom

## Abstract

**Purpose:** To investigate the role of hsa-miR-483-3p on the regulation of extracellular matrix (ECM) in cultured human trabecular meshwork (HTM) cells with known steroid response.

**Methods:** Primary cultures of HTM cells with known steroid responsiveness [GC-responder (GC-R) and GC-Non-responder (GC-NR) cells] were grown on coverslip in 12-well plate until 80% confluence and treated with 100 nM dexamethasone (DEX) for 24 h and transfected with different concentrations of synthetic miRNA 483-3p mimic or inhibitor. After 24 h or 72 h post transfection, the cells were harvest for the following experiments: (i) percentage transfection efficiency (ii) RNA isolation for qPCR analysis and (iii) immunofluorescence staining, (iv) protein isolation for Western blotting respectively. All experiments were performed in triplicate with three biological samples (n□=□3).

**Results:** GC-R HTM cells showed significantly higher expression of SMAD4 as compared to GC-NR HTM cells. Similarly, DEX treatment up-regulated the SMAD4-dependent ECM proteins. The presence of a miR-483-3p mimic down-regulated SMAD4 expression and SMAD4-dependent ECM production in a dose-dependent manner by negatively down- regulating SMAD4/TGF-β2 signalling. The inhibition of SMAD4-dependent ECM production by miR-483-3p was more pronounced in GC-R HTM cells as compared to GC- NR cells.

**Conclusion:** The down-regulation in ECM production by miR-483-3p is SMAD4 dependent and may play a protective role in mitigating steroid response in GC-R HTM cells. Using miR-483-3p mimics demonstrates therapeutic potential for the management of steroid induced ocular hypertension and glaucoma.

## Introduction

In the eye, the intraocular pressure (IOP) is generated and maintained as a balance between the aqueous humour formation and drainage through an architecturally complex tissue, the conventional/trabecular pathway. The trabecular meshwork (TM) at the irido-corneal angle forms the conventional outflow pathway which is the primary site for the aqueous humour to exit from the anterior chamber of the eye [1–3]. The TM outflow pathway provides resistance to the aqueous humour outflow and the majority of the resistance resides in the inner wall region of TM, which comprises the juxtacanalicular connective tissue (JCT) and the inner wall endothelium of Schlemm’s canal (SC) [4]. Dysregulation of this outflow pathway results in elevated IOP in glaucoma patients [5] and in susceptible individuals treated with steroids [6,7]. Glucocorticoids (GC) are steroids mainly prescribed for the management of ocular inflammatory and auto-immune diseases [8]. Topical steroids are being used to treat anterior segment inflammatory conditions whereas intravitreal steroids are recommended to treat posterior segment inflammatory eye diseases such as diabetic macular edema, non-infectious uveitis and retinal vein occlusion [9–11]. Long-term use of these steroids can cause elevated intraocular pressure (IOP). Topical treatment with steroids can raise IOP in about 30-40% of the individuals in normal population [12,13]. Intravitreal steroids use can cause moderate to severe IOP elevation and hence it becomes a major clinical challenge [14–16]. If left untreated, it can lead to secondary open angle glaucoma [17,18]. However, the molecular mechanism responsible for the differential steroid responsiveness is poorly understood.

In our previous study, we identified a unique miRNA signature between GC-responder (GC- R) and non-responder (GC-NR) cultured HTM cells using small RNA sequencing and it was found that hsa-miR-483-3p was up-regulated in both GC-R and GC-NR HTM cells [19]. HTM cells with known GC responsiveness were isolated from the GC characterized human donor eyes using an SI-OHT ex vivo model system (Haribalaganesh et al., 2021)Upregulation of miRNA-483-3p has been reported in diabetes and cardiovascular diseases and has been associated with cell apoptosis and lipotoxicity across diverse cell types [20]. In HTM cells, miR-483-3p was down-regulated under oxidative stress and over-expression of this miRNA had an inhibitory effect on extracellular matrix (ECM) production by inactivating SMAD4/TGFβ2 signaling pathway [21]. Therefore, in the present study the functional role of miR-483-3p on the regulation of ECM production in both GC-R and GC-NR HTM cells was also investigated.

Herein, we report that the expression of miR-483-3p and SMAD4 is inversely correlated in GC-R HTM cells. Overexpression of miR-483-3p negatively down-regulated SMAD4 expression in GC-R HTM cells and resulted in the decreased expression of SMAD4- dependent ECM proteins in GC-R TM cells. These results clearly indicate that miR-483-3p plays a protective role in mitigating the steroid response by down-regulating SMAD4 in GC- R HTM cells as compared to GC-NR HTM cells. Hence, miR-483-3p may have a therapeutic potential for the treatment of steroid-induced ocular hypertension (SI-OHT) and glaucoma (SIG).

## Materials and Methods

### Ethical Statement

Human donor eyes not suitable for corneal transplantation due to insufficient corneal endothelial cell count were used to establish the primary cultures of HTM cells as included in this study. The written consent of the deceased donor who were earlier consented to donate eyes after the death or next of kin was also obtained. The study protocol was approved by the Institutional Review Board of the Aravind Medical Research Foundation **(ID NO. RES2017006BAS)** and was conducted in accordance with the tenets of the Declaration of Helsinki.

### Human Donor Eyes

Human cadaveric eyes were obtained from the Rotary Aravind International Eye Bank, Aravind Eye Hospital, Madurai, India. The cadaveric eyes tissues were processed in accordance with the Declaration of Helsinki with approval of Institute Human Ethics Committee. The cadaveric eyes were enucleated within 5 h of death (mean elapsed time between death and enucleation was 3.60 ±0.99 h) and preserved at 4° C in the moist chamber until culture. The donor eyes used for this study is summarized in Supplementary Table S1.

In a set of paired eyes, one eye was used to establish a HOCAS ex vivo model system to characterize GC responsiveness after DEX treatment and the other eye was used to establish primary HTM cultures from eyes with identified responsiveness [22] (Supplementary Figure S1). The characteristics of donor eyes used for this study are summarized in Supplementary Table S1.

### Cell Culture and miRNA Transfection

Primary cultures of human trabecular meshwork cells with known steroid responsiveness were established as described previously [19,23]. GC-R and GC-NR HTM cells were cultured at 37 °C in 5% CO_2_ in low glucose Dulbecco’s Modified Eagle Medium (DMEM) with 15% fetal bovine serum, 5 ng/mL basic fibroblast growth factor and antibiotics. They were grown on coverslip in 12-well plate till 80% confluence and treated with 100 nM dexamethasone (DEX) for 24 h and transfected with different concentrations of synthetic miRNA 483-3p mimic or inhibitor or transfection negative control (Qiagen, MD, USA) using HiPerfect (Qiagen, Hilden, Germany) as per the manufacturer’s instructions. After 24 h or 72 h post transfection, the cells were harvest for the following experiments: (i) percentage transfection efficiency (ii) RNA isolation for qPCR analysis and (iii) immunofluorescence staining, (iv) protein isolation for Western blotting respectively. All experiments were performed in triplicate with three biological samples (nLJ=LJ3).

### Transfection Efficiency and Cell Viability

Primary cultures of GC-R and GC-NR HTM cells were transfected with synthetic miRNA mimic negative control with FAM label were transfected after 24 h DEX treatment using HiPerfect. After 24 h post-transfection, the HTM cells were fixed and observed under confocal microscopy and counted the cells showing FAM positivity. The percentage of transfection efficiency was calculated by using the following formula = No. of transfection- positive cells / Total no. of cells (DAPI) X 100. The cell viability was also assessed by tryphan blue dye exclusion assay.

### *Real-Time quantitative PCR (*RT-qPCR)

Total RNA was isolated from GC-R and GC-NR HTM cells 24 h post-transfection with respective treatments (miR-483-3p mimic or inhibitor or negative control) using Trizol reagent (Sigma, St. Louis, MO, USA) as per the manufacturer’s instructions. RNA quantity and quality were assessed by NanoDrop lite spectrophotometer (Thermofisher Scientific, Delaware, USA), First-strand cDNA was synthesized from total RNA using Verso cDNA kit (Thermo Scientific, Vilnius, Lithuania) according to the manufacturer’s instruction. Real- time PCR was performed with QuantiNova SYBR green (Qiagen, Hilden, Germany) in ABI- QuantStudio 5 (Applied Biosystems, Waltham, MA, USA). All mRNA were measured at CT threshold levels and normalized with the average CT values of a reference control (ACTB).

Values were expressed as fold increase over the corresponding values for control by the 2^− ΔΔCT^ method as described previously [23]. The list of primers of the selected genes for qPCR is shown in the Supplementary Table S2.

### Western Blot Analysis

Total protein was isolated from the transfected cells, the cells using Laemmli buffer and protease inhibitor cocktail (Tocris, Bristol, UK) after washing with ice cold PBS (Gibco, Thermo Scientific). The concentration of the isolated protein was estimated using BCA Protein Assay kit (Pierce, Thermo Scientific). Equal amount of protein (20 μg) from each sample was separated by 10% SDS-PAGE followed by transfer of resolved proteins onto nitrocellulose membrane. Membrane was blocked with 5% nonfat dry milk and incubated with primary antibodies [SMAD4 (1:1000), Abcam (Cat# ab110175); TGFβ2 (1:500), Abcam (Cat# ab167655); collagen 1A (1:250), Santa Cruz (Cat# sc-59772); fibronectin (1:100), Santa Cruz (Cat# sc-52331), laminin5 (1:500), Santa Cruz (Cat# sc-13587)] overnight at 4°C. Then the membrane was washed twice with TBS-T and TBS followed by incubation with secondary antibodies [anti-mouse IgG-HRP (1:1000), Abcam (cat# ab205719); goat anti-rabbit IgG-HRP (1:1000), Abcam (Cat# ab205718) for 1 h, followed by washing twice with TBS-T and TBS. The protein bands were detected using enhanced chemiluminescence Western Blotting Substrate (Thermo Scientific, IL, USA). Densitometry on immunoblot films was performed using NIH Image J sofware (http://imagei.nih.gov.ij/) provided in the public domain by NIH, Bethesda, MD, USA). Data were normalized to the loading control (anti-β–actin). Fold change in expression was calculated after normalization with control.

### Immunofluorescence analysis

The transfected GC-R and GC-NR HTM cells were washed twice with 1x PBS for 10 min and fixed with 4% paraformaldehyde for 30 min followed by washed with 1x PBS for 10 min. The fixed HTM cells were permeabilized using 0.2% Triton-X100 in PBS for 10 min. The endogenous biotin was blocked using avidin –biotin blocking system (Invitrogen, MD, USA) for 10 min. Further, the cells were incubated with primary antibodies [SMAD4 (1:500) (Cat# sc-7966); TGFβ2 (1:500) Cat# sc-374659); collagen 1A (1:250), Cat# sc-59772); fibronectin (1:200), Cat# sc-52331); laminin5 (1:500), Cat# sc-13587) all from Biotechnology, TX, USA] overnight at 4°C. After extensive washing with PBS, secondary antibody [anti-mouse IgG (1:300), Santa Cruz Biotechnology (cat# sc-516142), TX, USA] for 1 hr. They were then incubated with Dy-light 594 labelled streptavidin for 1hr, washed and mounted with anti-fade mounting media containing DAPI (Vector Labs, Inc. CA, USA). Images were captured using confocal microscope (Leica SP8 Confocal Microscope, Leica, Wetzlar, Germany). Cells without primary antibody served as a negative control.

### Statistical Analysis

Statistical analysis was carried out using Graph Pad Prism (ver.8.0.2) (Graph Pad software, CA, USA). All data are presented as mean ± SEM or otherwise specified. Statistical significance between two groups was analyzed using unpaired 2-tailed Student’s t test. P <0.05 or less was considered as statistically significant.

## Results

The transfection efficiency was more than 90% (Supplementary Figure S1) and the cell viability was more than 95 %.

### MiR-483-3p down regulates SMAD and TGFβ2 Signalling

Given SMAD4 is the direct target of miR483-3p as previously described, we examined whether miR483-3p targets SMAD4 in HTM cells after DEX treatment. Quantitative PCR and Western blot analysis revealed that DEX treatment significantly up-regulated the expression of SMAD4 in GC-R HTM cells [10 fold (qPCR) and (3 fold (WB)] as compared to GC-NR HTM cells. The presence of 483-3p mimic down-regulated the SMAD4 expression in a dose-dependent manner [5nM: 5 fold; 10nM: 10 fold and 25nM: 33 fold (qPCR); 5nM: 4 fold; 10nM: 5.5 fold and 25nM: 11 fold (WB)]. The presence of 483-3p inhibitor showed up-regulation of SMAD4 expression (Figure 1 & 2). Immunofluorescence analysis also revealed significantly higher expression of SMAD4 in GC-R HTM cells (Figure 3a).

**Figure 1.**
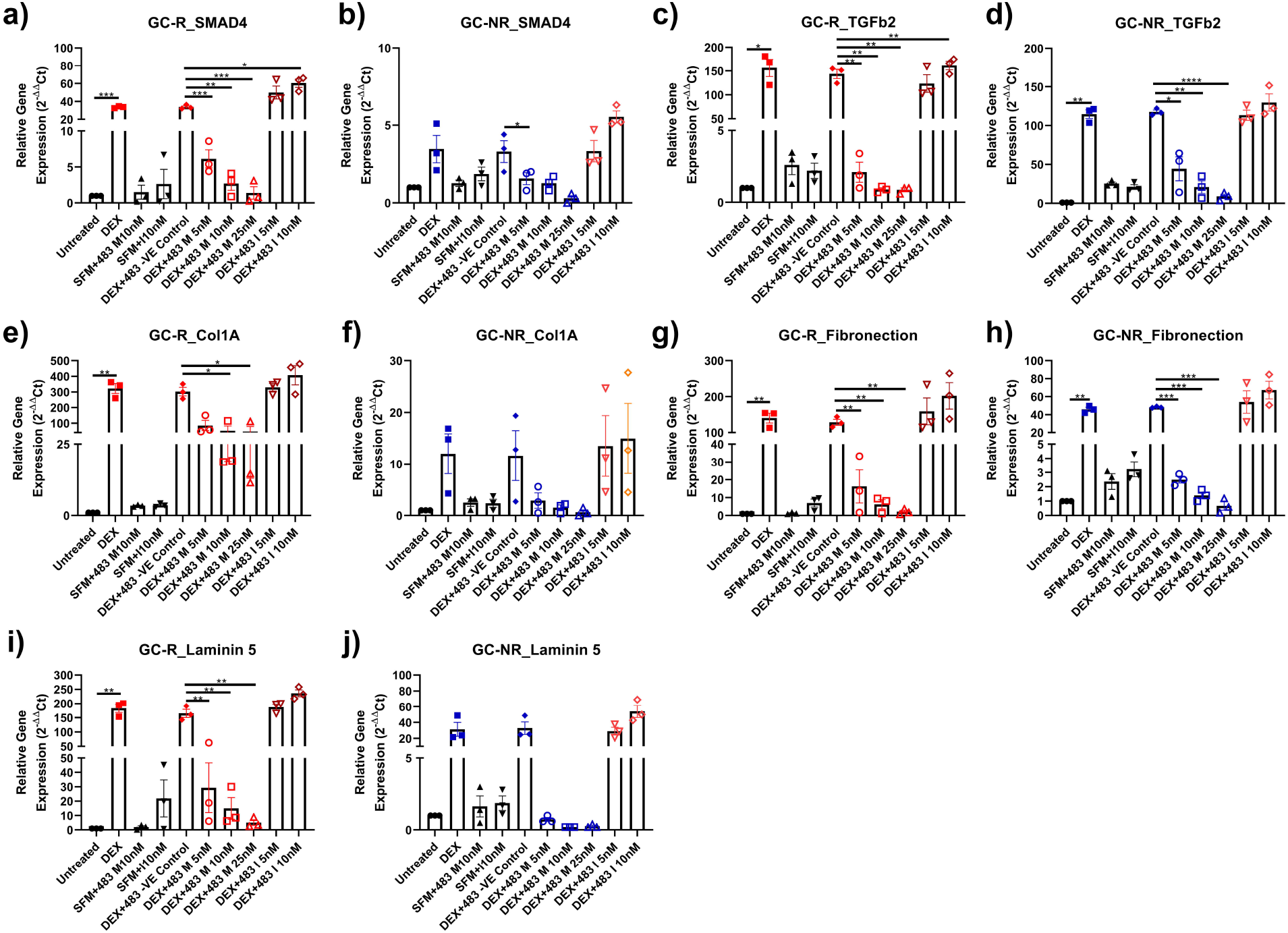
Effect of miR-483-3p by qPCR Analysis. HTM cells were transfected with hsa- miR-483-3p mimic, mimic negative control and inhibitor after 100nM DEX treatment. Total RNA was extracted, converted to cDNA and the SMAD4,TGFβ2,Collagen 1A, Fibronectin and Laminin5 gene expression were carried out by qPCR, gene expression were normalized to ACTB and analyzed using the 2^−ΔΔCT^ method. SMAD4- GC-R (a) GC-NR (b) HTM cells (n=3); TGFβ2 - GC-R (c) GC-NR (d) HTM cells (n=3); Collagen 1A GC-R (e) GC-NR (f) HTM cells (n=3); Fibronectin GC-R (g) GC-NR (h) HTM cells (n=3); Laminin5 GC-R (i) GC-NR (j) HTM cells (n=3). Data represent mean ± SEM; *p <0.05. **p <0.001; ***p <0.0001.

**Figure 2.**
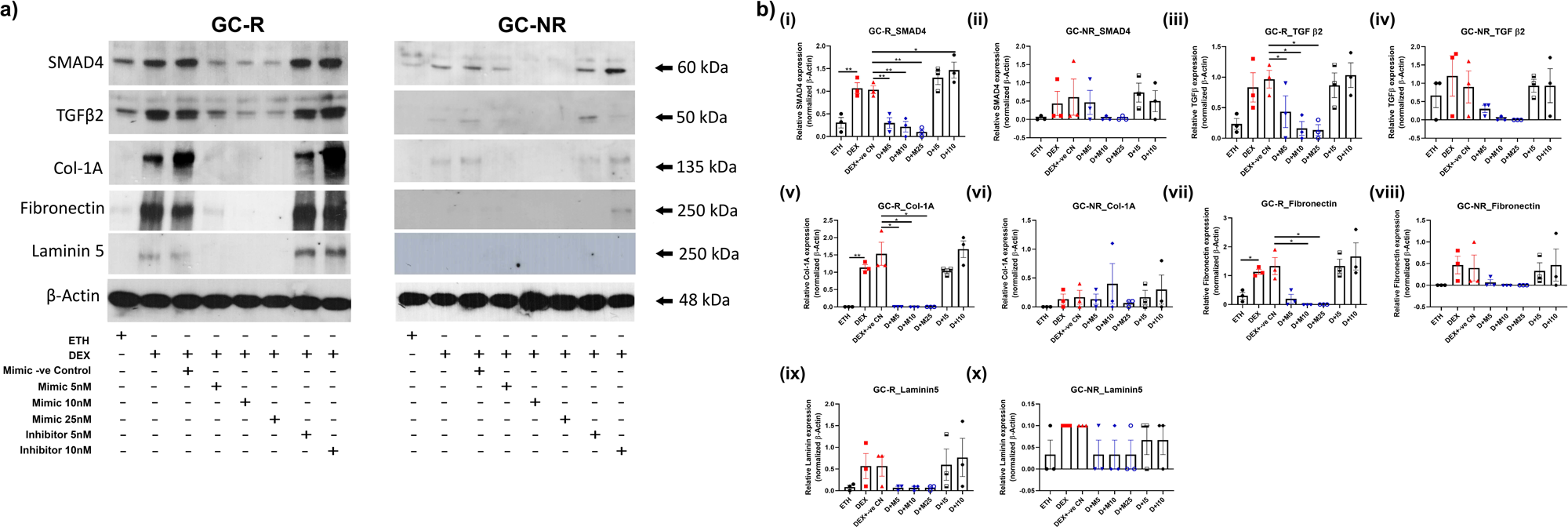
Western blot analysis of miR-483-3p target protein expression. miR-483-3p significantly downregulated SMAD4, TGFβ2 and ECM proteins in HTM cells (a). Western blot band intensity was used to measure differences in miR-483-3p target protein expression in GC-R and GC-NR HTM cells (b).

**Figure 3.**
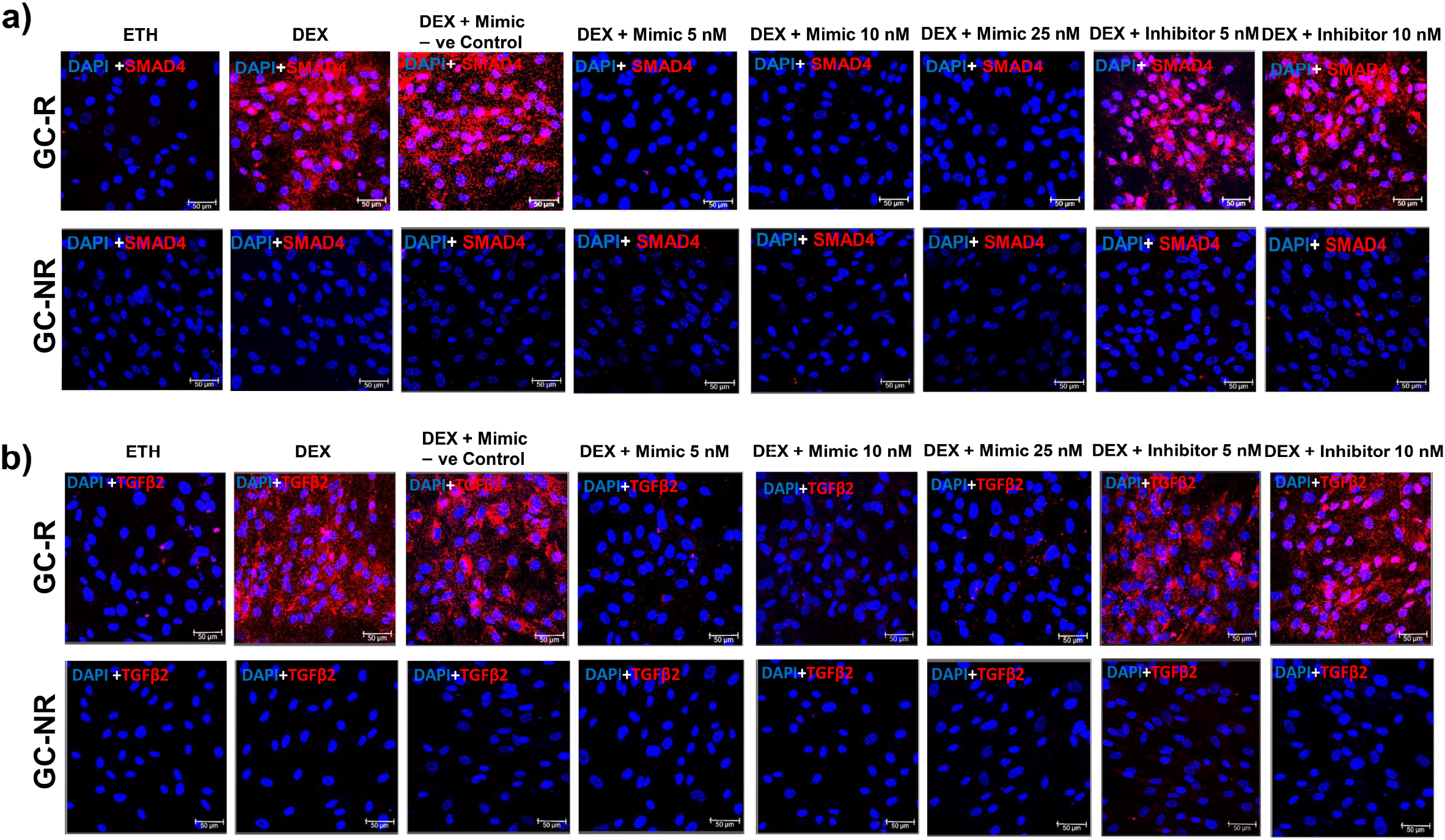
MiRNA-483-3p downregulates SMAD4 and TGFβ2 expression in HTM Cells. Representative confocal image of SMAD4 (a) and TGFβ2 (b) protein expression in hsa-miR- 483-3p mimic, negative control and inhibitor transfected GC-R and GC-NR HTM Cells.

**Figure 3c.**
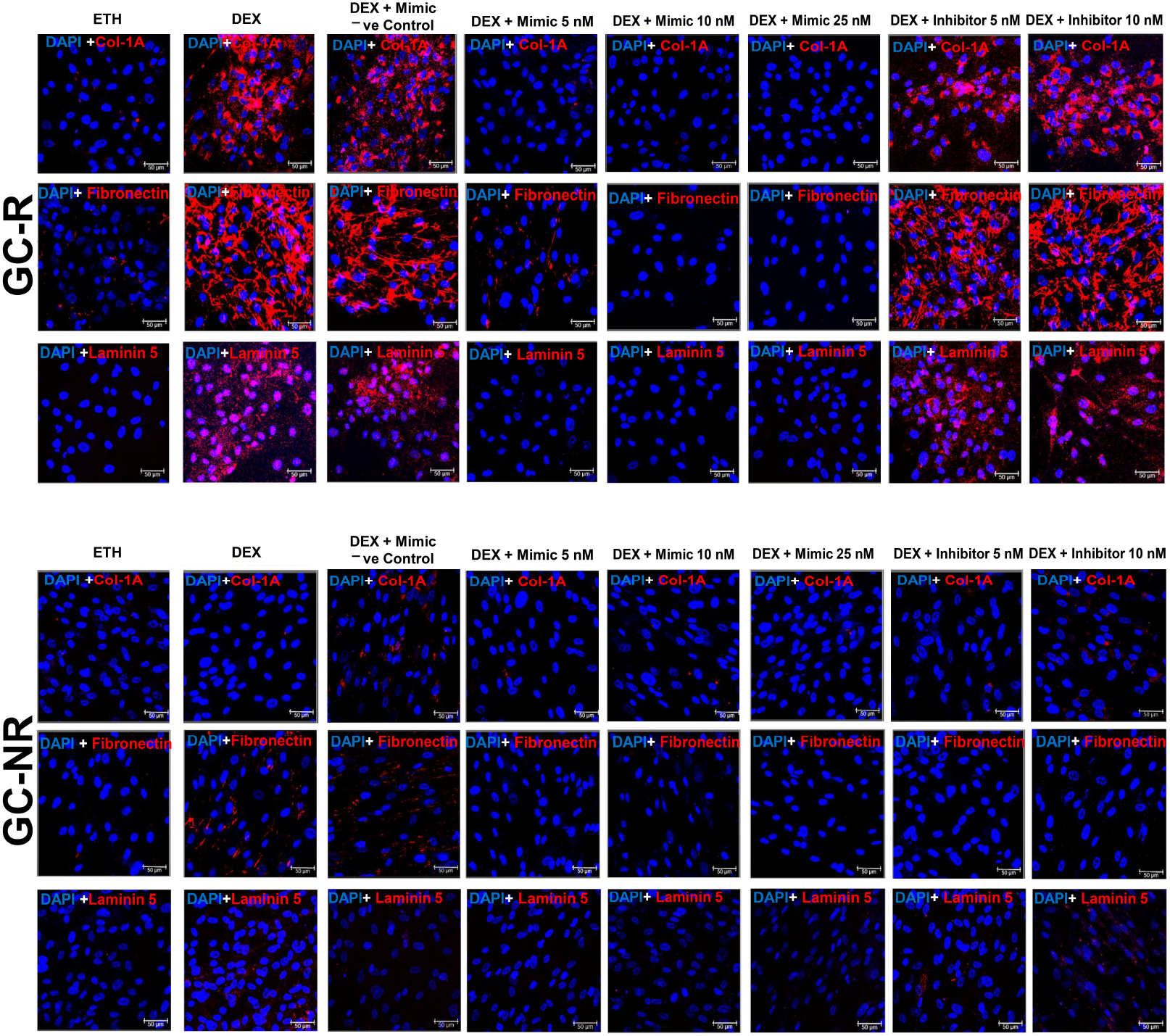
MiRNA-483-3p downregulates Collagen 1A, Fibronectin and Laminin 5 expression in HTMCs. Representative confocal image of Collagen 1A, Fibronectin and Laminin 5 protein expression in hsa-miR-483-3p mimic, negative control and inhibitor transfected GC-R and GC-NR HTM Cells.

Interestingly, the presence of miRNA 483-3p mimic negatively down-regulated SMAD4 expression and the effect is more prominent in GC-R HTM cells as compared to GC-NR cells. These results indicate that 483-3p targets SMAD4 and its expression was modulated by the presence of mimic and inhibitor (Figure 3a).

Similarly DEX treatment elevated SMAD4 dependent TGFβ2 expression in both GC-R and GC-NR HTM cells. The presence of 483-3p mimic down-regulated TGFβ2 expression in a dose-dependent manner. The presence of 483-3p inhibitor showed up-regulation of TGFβ2 expression in both GC-R and GC-NR by western blot and immunofluorescence analyses (Figure 2a & Figure 3b).

### MiR-483-3p down regulates SMAD4-dependent ECM proteins

In HTM cells, DEX treatment is known to induce the expression of ECM proteins as described previously [24]. Therefore, the effect of miR483-3p on ECM regulation was also studied in GC-R and GC-NR HTM cells. DEX treatment increased the expression of COL1A, fibronectin and laminin5 which was down-regulated by the presence of miR483-3p mimic in both GC-R and GC-NR cells by Western blot analysis (Figure 2a & Figure 3c). RT-qPCR and immunofluorescence analyses were in agreement with the Western blot analysis (Figure 2b v-x; Figure 3c). Interestingly, the down-regulation of ECM proteins by miR483-3p was more pronounced in GC-R and GC-NR HTM cells.

## Discussion

MicroRNAs (miRNAs) belong to a class of small non-coding RNAs that regulate complementary mRNA translation in eukaryotic organisms [25]. They were found to have a critical role in human aqueous humour production, absorption, maintenance of TM homeostasis, regulation of retinal ganglion cells apoptosis and glaucoma [25–27]. It is known to be one of the epigenetic drivers in glaucoma which strongly suggesting their involvement in the occurrence and development of glaucoma by regulating several biological processes of glaucoma-associated genes [28]. Several miRNAs were identified in glaucoma affected tissues/fluids such as aqueous humor, tears, TM and retina of patients with glaucoma and animal models [27,29–32].

To support the function of IOP homeostasis, several extracellular (ECM) proteins and their associated degradation enzymes were secreted by TM cells which are responsible for the continual remodelling of ECM [33]. Elevated IOP associated with steroid use is due to the remodeling of the TM tissue which causes an imbalance between the deposition and destruction of ECM in the TM cells which results in an increase in resistance to the aque`ous outflow` [33]. In the TM, steroid use induces structural, morphological and biochemical alterations which results in an increase in resistance to aqueous outflow and elevated IOP [33]. Following intra-vitreal injection of triamcinolone (IVTA) there is an up-regulation of proteins and miRNAs that are associated with ECM remodeling, cytoskeletal re-arrangement and mitochondrial oxido-reduction in IVTA-treated eyes of non-human primates [34]. IVTA up-regulated miR-29b and down-regulated miR-335-5p which are involved in oxidative stress and mitophagy whereas the up-regulated miR-15/16 cluster is involved in cell apoptosis in the TM. All these miRNAs contributed to GC-induced biochemical changes in TM and increased the outflow resistance post-IVTA injections in non-primates [34].

In POAG and SIG, phenotypic changes such as cross-linked actin networks (CLANs), abnormal deposition of ECM and decrease in phagocytosis are responsible for the restricted movement of aqueous outflow and hence elevated IOP [24]. In this context, integrins, mechanosensitive receptors were implicated in the glaucomatous phenotypic changes that regulate the cytoskeletal events essential for the maintenance of ECM formation, contractility and phagocytosis. Dysregulation of integrin signaling particularly αvβ3 integrin has been associated with several diseases including glaucoma. It has been noted by Farali et al. [35] that elevated expression of αvβ3 integrins was observed in human HTM cells upto 2 weeks even after DEX removal emphasizing the importance of integrins in the regulation of IOP [36]. In support of this observation, it is shown that in absence of exogenous DEX treatment induced glucocorticoid-induced matrix which triggered POAG-associated non-Smad-TGFβ2 signaling and also these changes were found to be correlated with aberrant changes in target ECM genes and proteins in HTM cells [24].

MicroRNA-483 is an anti-fibrotic miRNA [37]. Both the mature and passenger strands are biologically active and are known to target the 3’UTR of important fibrotic targets such as CTGF, PDGF-β, TIMP-2 and Bcl-2 [38–41]. In HTM cells, miR-483-3p was down-regulated under oxidative stress and over-expression of this miRNA has an inhibitory effect on ECM production by inactivating SMAD4/TGFβ2 signaling pathway [21]. Therefore, in the present study the functional role of miR-483-3p on the regulation of ECM production in both GC-R and GC-NR HTM cells was investigated. The activation of TGFβ/Smad4 pathway regulates ECM production in the fibrotic process of various diseases including glaucoma [42–44]. TGFβ2 is a major regulator of the ECM in the TM [45]. Elevated levels of TGFβ2 are found in the aqueous humor of patients with POAG and regulate the deposition of key ECM components such as fibronectin, collagen and laminin in TM which elevates IOP [2,46]. TGFβ2 is also over- expressed in dexamethasone induction in cultured HTM cells [24].

MiR-483-3p was up-regulated in several cancers such as colorectal, breast and pancreatic cancers [47]. SMAD4 was initially identified as a direct target for miR-483-3p in pancreatic cancer [39]. The expression of miR-483-3p and SMAD4 was inversely associated in pancreatic ductal adenocarcinoma tissues which indicates the ongogenic function of miR- 483-3p in the early pathogenesis of pancreatic ductal adenocarcinoma [48]. In the present study, miR-483-3p was up-regulated in both GC-R and GC-NR HTM cells. Interestingly, SMAD4 expression was significantly higher (around 10-fold) in GC-R HTM cells as compared to GC-NR HTM cells. Increased SMAD4 expression in GC-R HTM cells proportionately up-regulated the expression of ECM proteins such as Col1A, fibronectin and laminin; however, the expression levels of TGFβ2 in both GC-R and GC-NR did not differ significantly which clearly indicates that miR-483-3p and SMAD4 expression is positively correlated in GC-responsive HTM cells in the present study.

Overexpression of miR-483-3p in GC-R HTM cells negatively down-regulated SMAD4 expression which further correlates with the decreased expression of ECM proteins as compared to GC-NR HTM cells. This observation is in contrary to the previous study by Shen et al. [21] in which an oxidative stress insult (H_2_O_2_) reduced the expression of miR-483- 3p in HTM cells. However, in their study which mirrors our findings overexpression of miR- 483-3p and knockdown of SMAD4 caused a significant reduction in ECM proteins [21]. These results clearly indicate that miR-483-3p play a protective role in mitigating the steroid response by down-regulating SMAD4 in GC-R HTM cells as compared to GC-NR HTM cells. Further studies are warranted to investigate why there is a differential expression of SMAD4 in GC-R and GC-NR HTM cells? Collectively, the present study revealed that GC-R HTM cells expresses more SMAD4 and hence SMAD-dependent significant production of ECM proteins. The overexpression of miR-483-3p inhibited the SMAD4-dependent ECM production by negatively down- regulating SMAD4/TGFβ2 signaling and hence the effect was more pronounced in GC-R HTM cells as compared to GC-NR cells (Figure 4). Therefore, this miRNA may be protective in regulating ECM proteins in GC-responsive cells and may have a therapeutic potential for the management of steroid-induced OHT/glaucoma and POAG.

**Figure 4.**
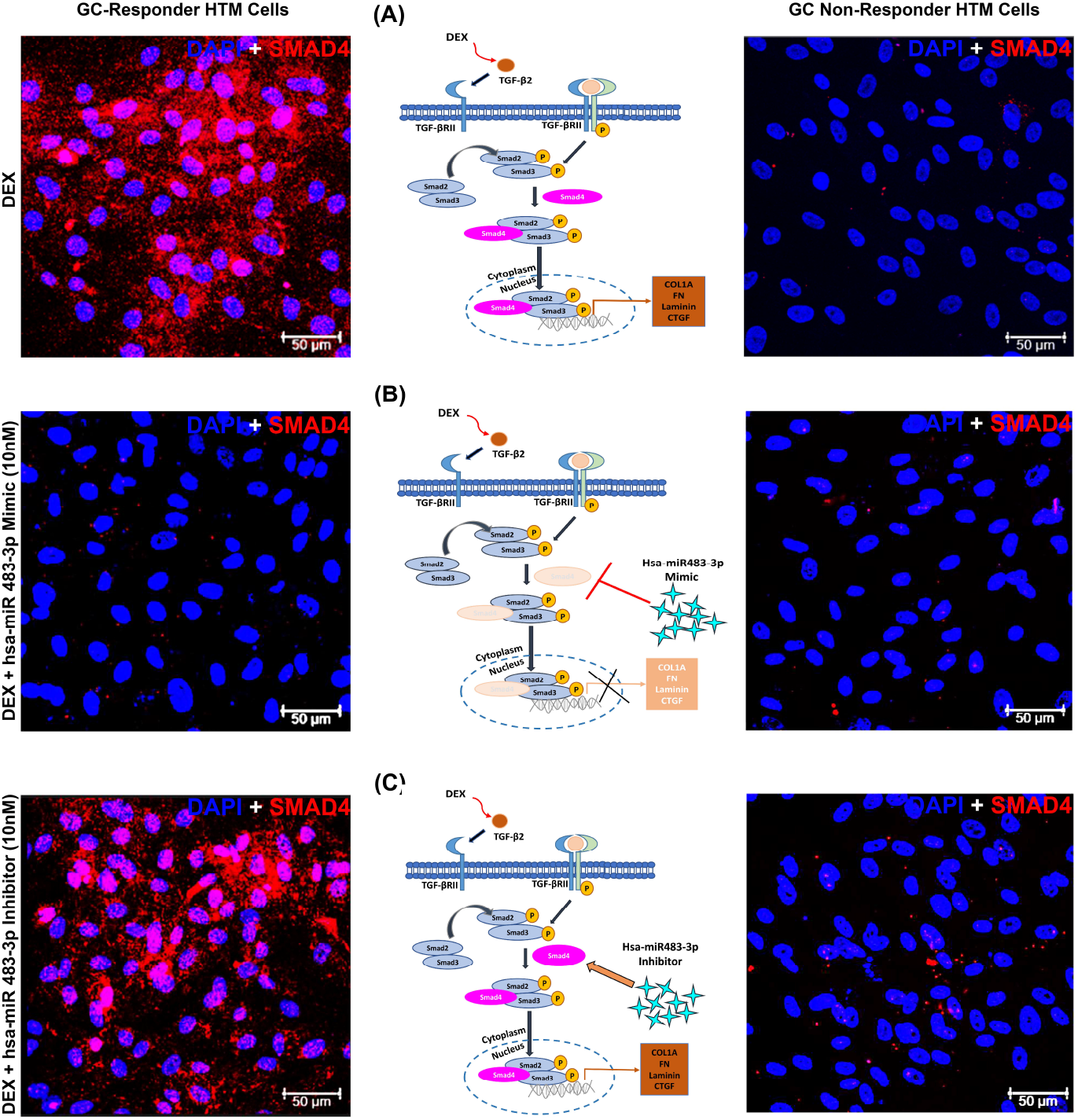
Proposed Schematic Representation of the Effect of miR483-3p on ECM Regulation in GC-R HTM cells. In GC-R HTM cells treatment with dexamethasone (DEX) activates canonical TGFβ2-SMAD signaling and causes dysregulated extracellular matrix (ECM) protein turnover resulting in the accumulation of ECM proteins which is implicated in steroid-induced ocular hypertension in GC-R HTM cells. The upregulation of TGFβ2-SMAD signaling by DEX (Panel A) in HTM cells. The upregulation of hsa-miR 483-3p inhibits the activation of TGFβ2-SMAD signaling by binding to SMAD4 mRNA leading to decreased SMAD4 protein (Panel B) in HTM cells. However, the presence of hsa-miR 483-3p antagomir/inhibitor decreases the expression of miR483-3p which in-turn enhances ECM deposition through the activation of TGFβ2-SMAD signaling (Panel C). Interestingly, the regulation of ECM by has-miR 483-3p was more pronounced in DEX-responder HTM cells (Left) as compared to DEX-non-responder cells (Right). TGFβR –TGFβ receptor complex; Col1A – Collagen 1A; FN – Fibronectin; CTGF-Connective tissue growth factor.

## Author Contributions

Conceptualization: S.S. and C.E.W.; Methodology: S.S. and C.E.W.; validation: S.S. and C.E.W.; formal analysis: R.H., S.S., and C.E.W.; investigation: R.H.; resources: S.S.; data curation: S.S. and C.E.W.; writing—original draft preparation: R.H., S.S., and C.E.W.; writing—review and editing: R.K. and R.S.; supervision: S.S. and C.E.W.; project administration, S.S.; funding acquisition: S.S. All authors have read and agreed to the published version of the manuscript.

## Institutional Review Board Statement

This study was approved by the standing Human Ethics Committee of Aravind Medical Research Foundation, Madurai, Tamilnadu, India (ID NO. RES2017006BAS).

## Informed Consent Statement

Not Applicable.

## Acknowledgments

The authors acknowledge the Rotary Aravind International Eye Bank, Aravind Eye Hospital, Madurai, India for providing human donor eyes for this study.

