## Supplementary file for "Hsa-MiR-483 -3p Regulates the Extracellular Matrix Proteins via TGFβ2/SMAD4 Signaling in the Glucocorticoid-responsive Human Trabecular Meshwork Cells"

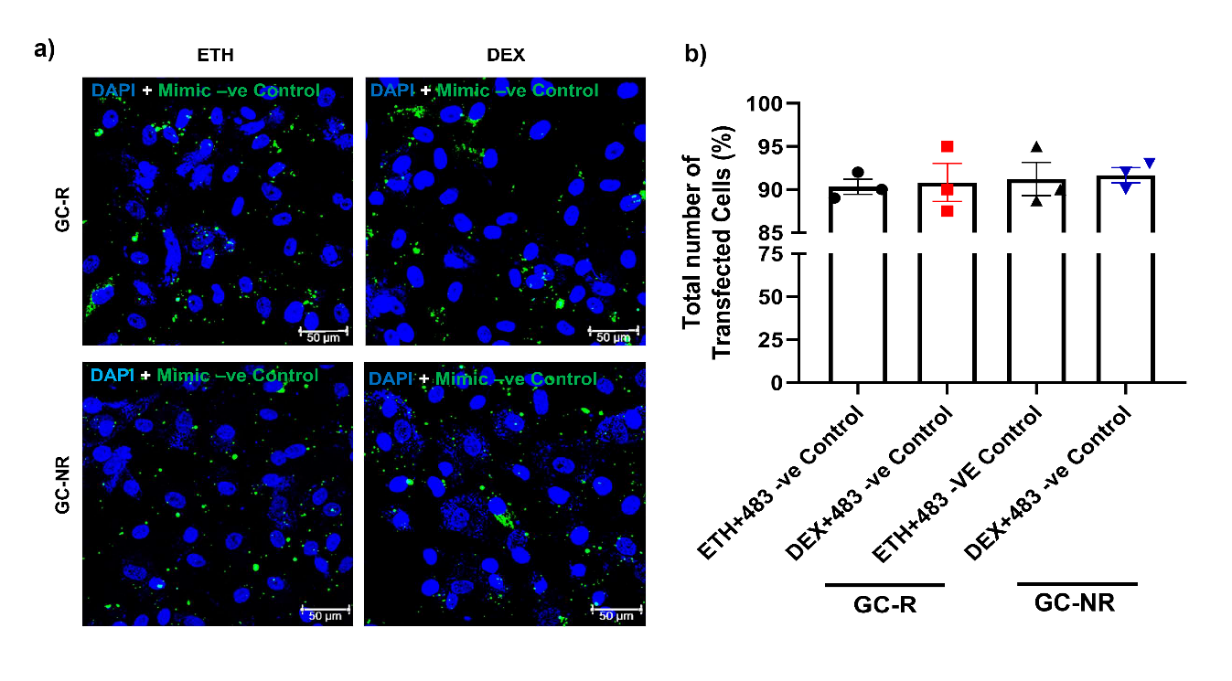

**Supplementary Figure S1.** Transfection efficiency analysis in primary HTM Cells. (a) Representative confocal image of hsa-miR-483-3p mimic negative control transfected in GC-R and GC-NR HTM cells with; (b)Graph showing the transfection percentage of GC-R (n=3) and GC-NR (n=3) HTM Cells.

**Supplementary Table S1:** Characteristics of Human Donor Eyes Used for the Study

| **Code** | **Age** | **Sex** | **Cause of Death** | **Time B/W Death & Enucleation (h)** | **Time B/W Enucleation & Culture (h)** | **Experiment**  **Eye** | **Treatment** | **GC R/NR** |
| --- | --- | --- | --- | --- | --- | --- | --- | --- |
| **TM Cells** | | | | | | | |  |
| OCHD19-01 | 73 | M | Natural | 4.5 | 46.91 | OS | ETH/DEX | NR |
| OCHD19-08 | 67 | M | Heart disease | 2.9 | 56 | OS | ETH/DEX | NR |
| OCHD20-10 | 38 | M | CKD | 1.41 | 47.5 | OS | ETH/DEX | NR |
| OCHD19-34 | 75 | F | Heart disease | 2.5 | 52.16 | OS | ETH/DEX | R |
| OCHD20-04 | 65 | M | COPD | 0.33 | 28.5 | OS | ETH/DEX | R |
| OCHD21-01 | 75 | M | Respiratory disease | 3.08 | 84.33 | OS | ETH/DEX | R |

^*^A total of 6 eyes were used for the study with the mean (±) SD age was 65.5±14.1 years. The mean (±) SD elapsed time between (b/w) death and enucleation was 2.45 ±1.4 h and the mean elapsed time between enucleation and culture was 52.57±18.2 h. b/w: between.

**Supplementary Table S2:** Primer pairs used for qPCR

| **S.No** | **List of Genes** | **Primer** |
| --- | --- | --- |
| 1 | SMAD4 | F 5'-CTACCAGCACTGCCAACTTTCC-3' |
|  |  | R 5'-CCTGATGCTATCTGCAACAGTCC-3' |
| 2 | TGFβ2 | F 5'-AAGAAGCGTGCTTTGGATGCGG-3' |
|  |  | R 5'-ATGCTCCAGCACAGAAGTTGGC-3' |
| 3 | Collagen 1A | F 5'-GATTCCCTGGACCTAAAGGTGC-3' |
|  |  | R 5'-AGCCTCTCCATCTTTGCCAGCA-3' |
| 4 | Fibronectin | F 5’-AATCCAAGCGGAGAGAGAGTCA-3' |
|  |  | R 5’-CATCCTCAGGGCTCGAGTAG-3' |
| 5 | Laminin5 | F 5'-GTCACAGAGCAGGAGGTGGCT-3' |
|  |  | R 5'-GCTTCTGTCAAGACTCTCCAGG-3' |
